# PrediRep: Modelling hierarchical predictive coding with an unsupervised deep learning network

**DOI:** 10.1101/2024.05.04.592511

**Authors:** Ibrahim C. Hashim, Mario Senden, Rainer Goebel

## Abstract

Hierarchical predictive coding (hPC) proposes that the cortex continuously generates predictions of incoming sensory stimuli. Deep neural networks inspired by hPC are frequently used to probe the neurocomputational mechanisms suggested by the theory in silico and to generate hypotheses for experimental investigations. However, these networks often deviate from hPC by prioritizing computational efficiency over alignment with its principles. To remedy this, we introduce PrediRep, a deep learning model explicitly designed to emphasize alignment with the theory. PrediRep incorporates the principles of hPC found in the other networks, while avoiding their deviations from it. We evaluate the performance of PrediRep on a next-frame prediction task and its functional alignment with hPC, comparing it to other contemporary deep learning networks inspired by the theory. Our findings demonstrate that PrediRep achieves the closest functional alignment with hierarchical predictive coding without sacrificing computational performance.

## 1. Introduction

The visual input that we receive from the world is intrinsically dynamic, subject to both internally and externally generated movements. Predicting future sensory input requires the cortex to surmise the latent physical causes, such as the shapes, positions, and speeds of objects that generate this sensory information. Hierarchical predictive coding (hPC) proposes that this can be achieved by minimizing prediction errors of an internal model representing the external world (Mumford, 1992; Rao and Ballard, 1998, 1999; Friston, 2003; Lee and Mumford, 2003; Friston, 2005; Huang and Rao, 2011; Clark, 2013; Jiang and Rao, 2022). Specifically, hPC postulates the existence of two distinct types of neurons in each cortical area, namely representation (R) and error (E) neurons. Representation neurons at one level of the cortical hierarchy are responsible for predicting the activities of R neurons at the immediate lower level of the hierarchy. At the lowest level, the R neurons predict the sensory input (Rao and Ballard, 1999). The predictions of future activities are enabled by local recurrent synapses of the R neurons (Rao and Ballard, 1998, 1999; Jiang and Rao, 2022). The E neurons compare the predicted activities of R neurons at the same hierarchical level with their actual activities, thereby computing prediction errors. These prediction errors are subsequently relayed to the R neurons at the same hierarchical level as well as to the level directly above (Rao and Ballard, 1999). The R neurons in the higher cortical areas utilize the prediction errors to refine their predictions. On the other hand, the R neurons at the same level leverage these errors to adjust their activity in line with the predictions and thus ensure consistency (Rao and Ballard, 1999; Spratling, 2017).

The algorithmic principles of hPC present a promising framework for facilitating unsupervised learning based on future state prediction, such as the prediction of subsequent frames in video sequences (Softky, 1995; Palm, 2012; O’Reilly et al., 2014; Goroshin et al., 2015; Srivastava et al., 2015; Mathieu et al., 2015; Patraucean et al., 2015; Finn et al., 2016; Vondrick et al., 2016; Lotter et al., 2016, 2020; Hosseini and Maida, 2020; Straka et al., 2023). This potential has previously been exploited by several implementations such as PredNet (Lotter et al., 2016, 2020), the Rao and Ballard Protocol (RBP; Hosseini and Maida, 2020), and PreCNet (Straka et al., 2023). These deep learning models, inspired by hPC, integrate hierarchically connected levels featuring both R and E units. Furthermore, they align with the proposed cortical goal of minimizing the activities of E units. These networks have since been leveraged to explore the implications of the hPC mechanism in the brain. PredNet, for instance, has been employed to investigate the role of hPC in human static illusions (Lotter et al., 2020) and motion illusions (Watanabe et al., 2018; Kirubeswaran and Storrs, 2023).

While PredNet, RBP and PreCNet do exhibit several fundamental similarities with hPC, these networks were nonetheless designed with a primary focus on next-frame prediction performance. As a result, certain deliberate design choices in the networks introduced disparities with principles of hPC. For instance, in PredNet, E unit activity is predicted, which contrasts with the premise of hPC that R unit activity should be predicted (Lotter et al., 2016). In RBP, the lower-level R unit activity is modified before being predicted, and therefore there is no prediction of immediate R unit activity as proposed by hPC (Hosseini and Maida, 2020). In PreCNet the activities of lower-level R units are directly predicted. However, the predictions are compared to the R unit activities for both the current and the future sensory input (Straka et al., 2023). This creates a multi-layered comparison process that deviates from hPC. Furthermore, in all three networks the R units are modeled as ConvLSTM units (Shi et al., 2015). ConvLSTM units are highly complex as they contain several intrinsic connections and operations. Using ConvLSTM units puts a strong emphasis on the learning within a level, whereas hPC postulates that a substantial portion of computational processing and learning should occur between levels (Rao and Ballard, 1999; Huang and Rao, 2011).

As previously discussed, the deviations of existing deep learning models from hPC stem from the algorithms being primarily tailored towards performance in predicting the next frame. We believe that in order to construct a model suitable for investigating hPC, the primary focus should shift to crafting a model that aligns as much as possible with the principles of hPC. We present PrediRep (PREDIcting REPresentations), an hPC-inspired deep learning network that adheres more closely to the hPC principles than any of the other networks. PrediRep achieves greater alignment with hPC by making two essential architectural choices. First, it incorporates hierarchical levels that directly predict the future activities of R units of lower levels. Second, PrediRep opts for the use of simple recurrent convolutions instead of complex ConvLSTM units for the recurrence of R units. We demonstrate that these implementation choices endow PrediRep with enhanced functional alignment with hPC. Remarkably, PrediRep also achieves competitive levels of performance on a next-frame prediction task while having significantly less learnable parameters. This marks a significant next step toward effectively integrating the hPC theory into a deep learning framework.

## 2. Methods

### 2.1. The PrediRep Model

PrediRep is composed of hierarchically stacked levels, each of which contains an R unit and an E unit (see Figure 1). The operations of PrediRep can be split into a downward and an upward sweep. During the downward sweep, information flows from higher to lower R units through feedback connections. They employ local recurrent connections to update their activity based on the feedback signal before transmitting their own activity further down the hierarchy. Simultaneously, through a second feedback connection, the R units generate predictions (Ậ), which are transmitted to the E units of the level below. After the downward sweep, an upward sweep occurs. During this phase, the E units calculate prediction errors and send their activity forward to the R units of the level above. These, in turn, update their activity again using local recurrent connections, this time based on the prediction errors. The updated activities of the R units are then sent through lateral connections to the E units of the same level in the form of targets (A). At the lowest level of the hierarchy, the target is the current input (I). Therefore, except for R1, the R units are engaged in predicting a mixture of their own past and current activities, refined by the input.

**Figure 1:**
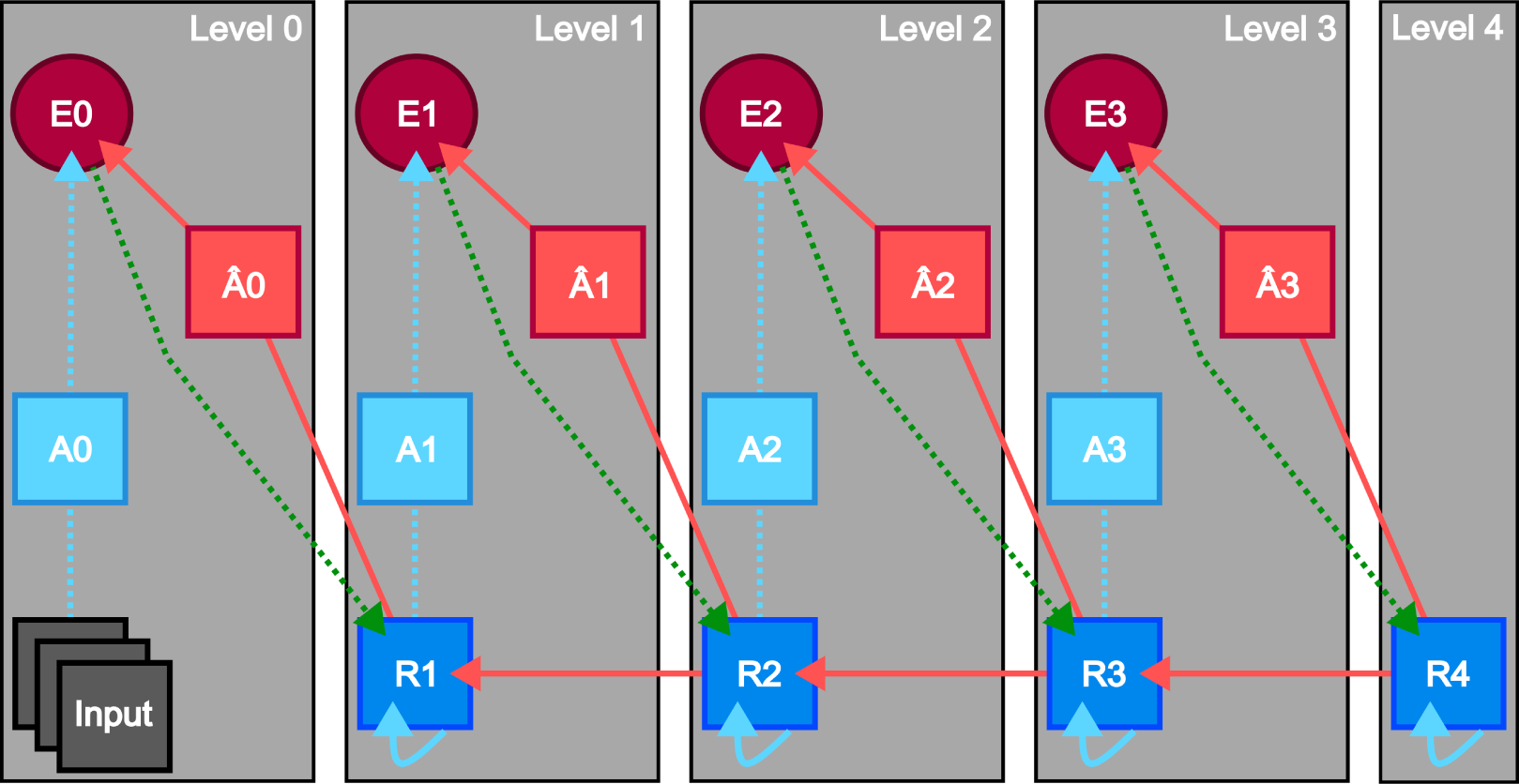
Visualization of the PrediRep Architecture. A detailed five-level implementation of PrediRep. E signifies error unit, R denotes representation unit, A represents target and A^^^ symbolizes prediction. Key connectivity patterns within the architecture are represented by combinations of line colors and styles. Red lines depict feedback connections, blue lines lateral connections and green lines feedforward connections. Solid lines indicate trainable connections and dotted lines non-trainable connections.

Algorithm 1 illustrates all operations used in PrediRep in order of their execution. Notably, the R units play distinct roles in the upward and downward sweeps. During the downward sweep, the R units predict the activity of a pooled version of their counterparts in the level below during the subsequent upward sweep. The pooling allows for convergence to occur between levels. The design of PrediRep ensures that the information from the input frame reaches all levels during the upward sweep. To achieve recurrences in the R units convolutions are employed over the concatenated inputs allowing the integration of information from various sources. PrediRep uses rectified linear units (ReLU) for the non-linearities of the R and E units. This means that the activities of the unit are forced to be positive, allowing for comparability with neuronal activity and aligning with hPC. Most predictions within PrediRep are also generated using a ReLU non-linearity. However, in the lowest level a Saturated Linear Unit (SatLU) is used with values ranging between 0 and 1. This choice aligns with the pixel value ranges in the input frames.

#### Algorithm 1

Calculation of States in Network. Terms are *F* (frames), *f* (current frame), *l* (level), *L* (final level), *Rdw* (representation layer downward sweep), *Rup* (representation layer upward sweep), *E* (error layer), *A* (target), Ậ (prediction).

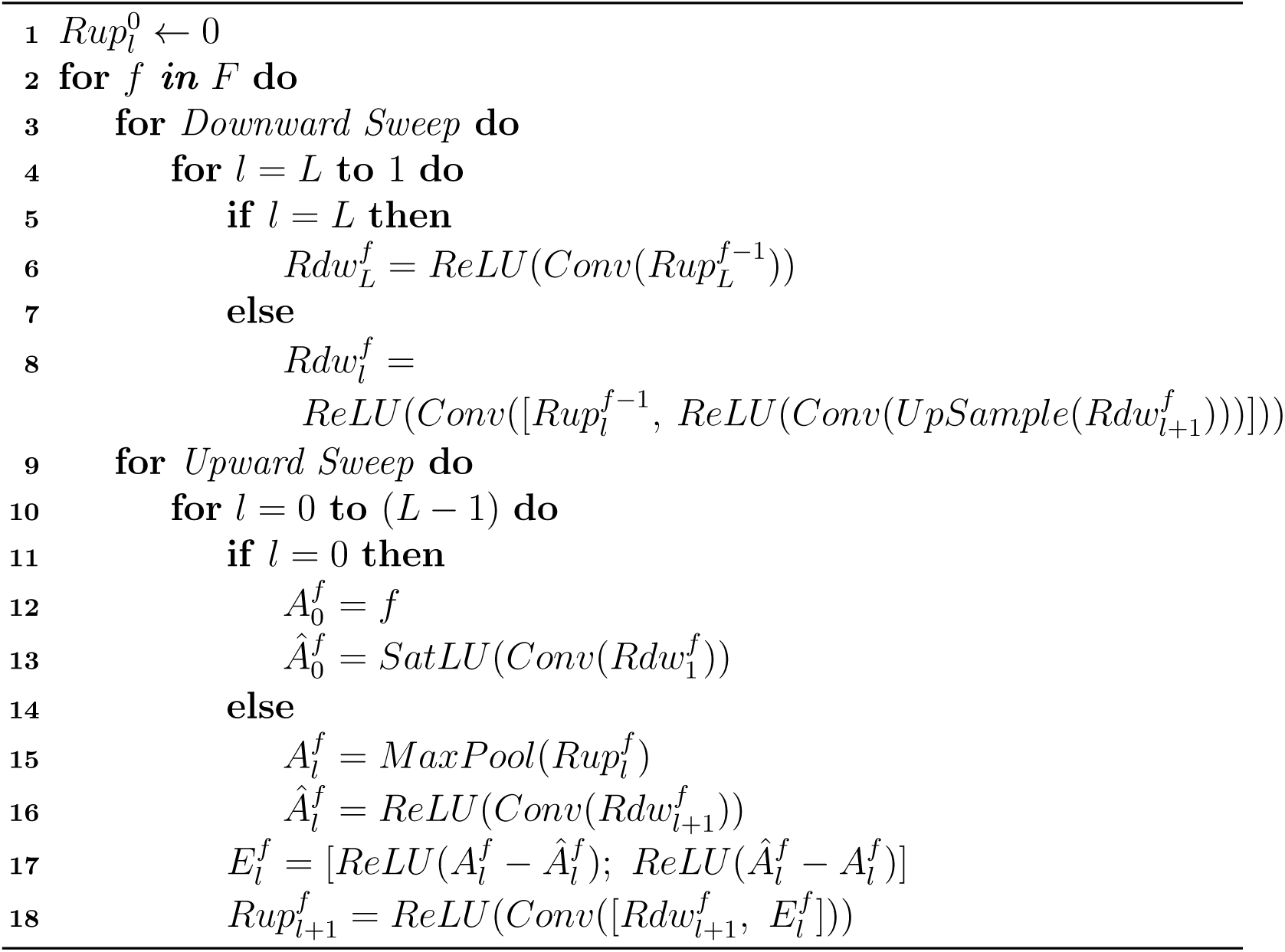

Equation 1 illustrates the loss function of PrediRep, which is computed as a sum of activities across the E units.

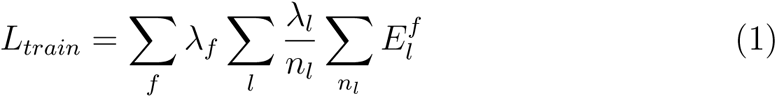

Their activities are aggregated across all presented frames and levels to calculate the overall loss. Within this equation, *λ_f_*signifies the weight allocated to each frame in the loss computation while *λ_l_* denotes the weight attributed to each of the units in the loss function. The *n_l_*term acts as a normalization factor, scaling the E unit activities by their respective sizes. The integration of information from Algorithm 1 and Equation 1 reveals the two operational stages of PrediRep, namely the inference and the learning stage. During the inference stage the R and E units are updated over several frames and the activities of the E units are recorded for each frame. Then, during the learning stage, the total activity of the E units across the frames is used as the loss signal to update the weights of the learnable connections in PrediRep.

### 2.2. Datasets

Two distinct video datasets were utilized, both capturing urban driving environments from the perspective of a car. These datasets are the KITTI datasets (Geiger et al., 2013) and the CalTech Pedestrian dataset (Dollar et al., 2009). Both datasets were utilized in previous works when investigating hPC-inspired deep learning networks (Hosseini and Maida, 2020; Lotter et al., 2016, 2020; Straka et al., 2023). The KITTI dataset was employed for training and testing, whereas the CalTech dataset was used only for testing. KITTI is a dataset comprising videos recorded at 10 frames per second captured from a roof-mounted camera on a car navigating urban regions in Germany. CalTech Pedestrian is a dataset containing videos obtained at a higher frame rate of 30 frames per second from a dashboard-mounted camera in a car travelling through Los Angeles. Both datasets were preprocessed as described in Lotter et al. (2016). For KITTI a train set consisting of 41396 frames, a validation set containing 154 frames and a test set comprising 832 frames were created from the City, Residential and Road categories. This ensured that a diverse representation of urban environments was included. Each frame was cropped and resized to have dimensions of 128×160 pixels. For CalTech Pedestrian, 121465 frames were extracted from the original test set. Similarly to the KITTI dataset, these frames were cropped and resized to 128×160 pixels. Additionally, the frame rate was downsampled by a factor of 3 to align with the frame rate of the KITTI dataset.

### 2.3. Network Parameters

A total of four distinct hPC-inspired deep learning architectures were trained and compared. These architectures consisted of PrediRep, PredNet, PreCNet and the Rao and Ballard Protocol. To the best of our knowledge these four networks are the only hPC-inspired deep learning networks incorporating R and E units together with the aim of minimizing E unit activity trained on video sequences. PrediRep was implemented as outlined in Section 2.1. PreCNet was implemented as described in Straka et al. (2023), PredNet as in Lotter et al. (2016, 2020), and RBP as in Hosseini and Maida (2020). Unlike PrediRep, PreCNet, PredNet and RBP have all been designed with ConvLSTM units. ConvLSTM units typically use a hyperbolic tangent as the output activation function, which in PredNet leads to better computational performances in terms of next-frame prediction than alternative non-linearities, such as ReLU. However, this also leads to a reduction in the biological interpretability of the units as their outputs are bound between –1 and 1 and hence not well aligned with the strictly positive firing rates of neurons (Lotter et al., 2020). We therefore took the same approach as Lotter et al. (2020) for testing the functional alignments of the networks with hPC and used versions of the PreCNet, PredNet and RBP networks configured with a ReLU non-linearity for the recurrent units. However, for the computational results we also evaluated the networks using a Tanh non-linearity for the recurrent units. PrediRep was not tested with a Tanh activation function as it does not contain ConvLSTM units and therefore would not benefit from it. All four networks were designed as five-level models, each consisting of four R and E units. The exact implementations of all the networks can be found in our GitHub repository.^1^.

In this study, we decided to use the same number of filters across all networks as different numbers of filters in specific units could affect the computational performances of the networks. Previously, Straka et al. (2023) demonstrated that PreCNet outperformed PredNet when the two networks shared similar parameter numbers. However, their PreCNet architecture featured a substantially higher number of filters in the lowest R unit compared to their PredNet implementation. We therefore opted for the R units of the networks having (8, 16, 32, 64) filters and the Ậ units having (3, 8, 16, 32) filters. This configuration culminated in 64 channels constituting the R unit in the final level of each network. We deliberately opted for small dimensions to facilitate the training and testing of a substantial number of iterations for each network. To maintain consistency and comparability, we kept all other parameters consistent with those implemented in Lotter et al. (2016). The convolution operations in the networks utilized a kernel size of (3 x 3) with padding set to “same”. Max pooling was employed as the pooling operation, and both the upsampling and pooling operations used a kernel size of (2 x 2). An overview of the number of trainable parameters per network is given in Table 1.

**Table 1:** Number of Trainable Parameters of the Networks.

| Network | Number of Trainable Parameters |
| --- | --- |
| PreCNet | 561,427 |
| RBP | 524,971 |
| PredNet | 524,827 |
| PrediRep | 244,131 |

### 2.4. Training Parameters

All networks were trained on a NVIDIA GeForce RTX 2080 Ti using TensorFlow 1.6. The training process of every network employed the loss function defined in Equation 1 together with Backpropagation Through Time (Werbos, 1990). There were three distinct loss types utilized. These loss types varied in their weighting of *λ_l_* for each of the units, as reported in Table 2. Zero loss focused solely on the lowest level E unit. All loss included all E units in the loss calculation but with an emphasis on the one from the lowest level. Finally, Equal loss distributed the weighting equally across all E units. Zero loss was utilized for investigating PredNet, PreCNet and RBP previously and All loss for PredNet and RBP (Lotter et al., 2016, 2020; Hosseini and Maida, 2020; Straka et al., 2023). Results with Equal loss have, to the best of our knowledge, not been shown before for these networks. The weighting of the *λ_f_* variable was consistent across the loss types. For the first frame it was set to zero as no accurate prediction could be made yet and for all subsequent frames it was set to an equal value so that each frame was equally important for the loss. All networks underwent training with the three loss types. Each combination of network and loss type was independently trained 30 times.

**Table 2:** Weighting of Error Units Depending on Loss Type.

| Loss Type | Weighting of E0 | Weighting of Other E Units |
| --- | --- | --- |
| Zero | 1.0 | 0.0 |
| All | 1.0 | 0.1 |
| Equal | 1.0 | 1.0 |

The training of each network closely followed the procedure outlined in (Lotter et al., 2016) to maintain consistency and comparability. Training was conducted over 150 epochs with batch sizes set to contain 4 sequences. For each epoch 500 random sequences were utilized in total. The same frame could appear in more than one of these sequences. For validation 100 unique sequences, different to those used in training, were employed, with each frame being restricted to appearing in only one sequence. The Adam optimizer (Kingma and Ba, 2014) was used for training with default initial hyperparameter values (*α* = 0.001, *β*_1_ = 0.9, *β*_2_ = 0.999). The learning rate began at 0.001 for the first 75 epochs and was reduced by a factor of 10 for the remaining epochs.

### 2.5. Testing next frame prediction capabilities of networks

To assess the next-frame prediction capabilities of the networks, evaluations were conducted on the test sets of both the KITTI and the CalTech Pedestrian datasets utilizing two key metrics. The metrics were the Mean Squared Error (MSE) and the Structural Similarity Index (SSIM; Wang et al., 2004). These metrics have been used in previous works investigating the networks (Lotter et al., 2016, 2020; Hosseini and Maida, 2020; Straka et al., 2023). The testing on KITTI was conducted to investigate how effectively the networks learned the specific environment in which they were trained. Meanwhile, the testing on CalTech Pedestrian was undertaken to ascertain the robustness of the representations formed (Lotter et al., 2016). The transition from a roof-mounted to a dashboard-mounted camera significantly changes the field of view, potentially strongly impacting the performances of the networks. Conversely, the shift between countries, although introducing some variability, is less likely to have a substantial effect as both urban scenes share common objects such as cars, pedestrians, and buildings. For each network instantiation, MSE and SSIM values were computed for sequences comprising ten frames across the entire test sets of both datasets. The frames were unique to each sequence. The significance of the differences in the performances of the networks was established through a Welch’s t-test with Bonferroni correction.

## 3. Results

### 3.1. Next Frame Prediction Performance

Figures 2 to 5 provide a comprehensive analysis of the effectiveness of the representations acquired by the networks in predicting the next frame in video sequences. In Figure 2 the performances of the networks trained with Zero loss are depicted. All networks used a Tanh as the activation functions in their recurrent units, whilst PrediRep used a ReLU. PredNetZeroTanh significantly outperformed all other networks for both the MSE and SSIM metric on both datasets. PrediRepZero demonstrated equal performances to PreCNetZeroTanh for all metrics except for the MSE result on the CalTech Pedestrian dataset, where it underperformed it. PrediRepZero outperformed RBPZeroTanh across all metrics. All the networks showed a small amount of variation, with PreCNetZeroTanh demonstrating the widest range of performances with a larger standard deviation compared to the other networks.

**Figure 2:**
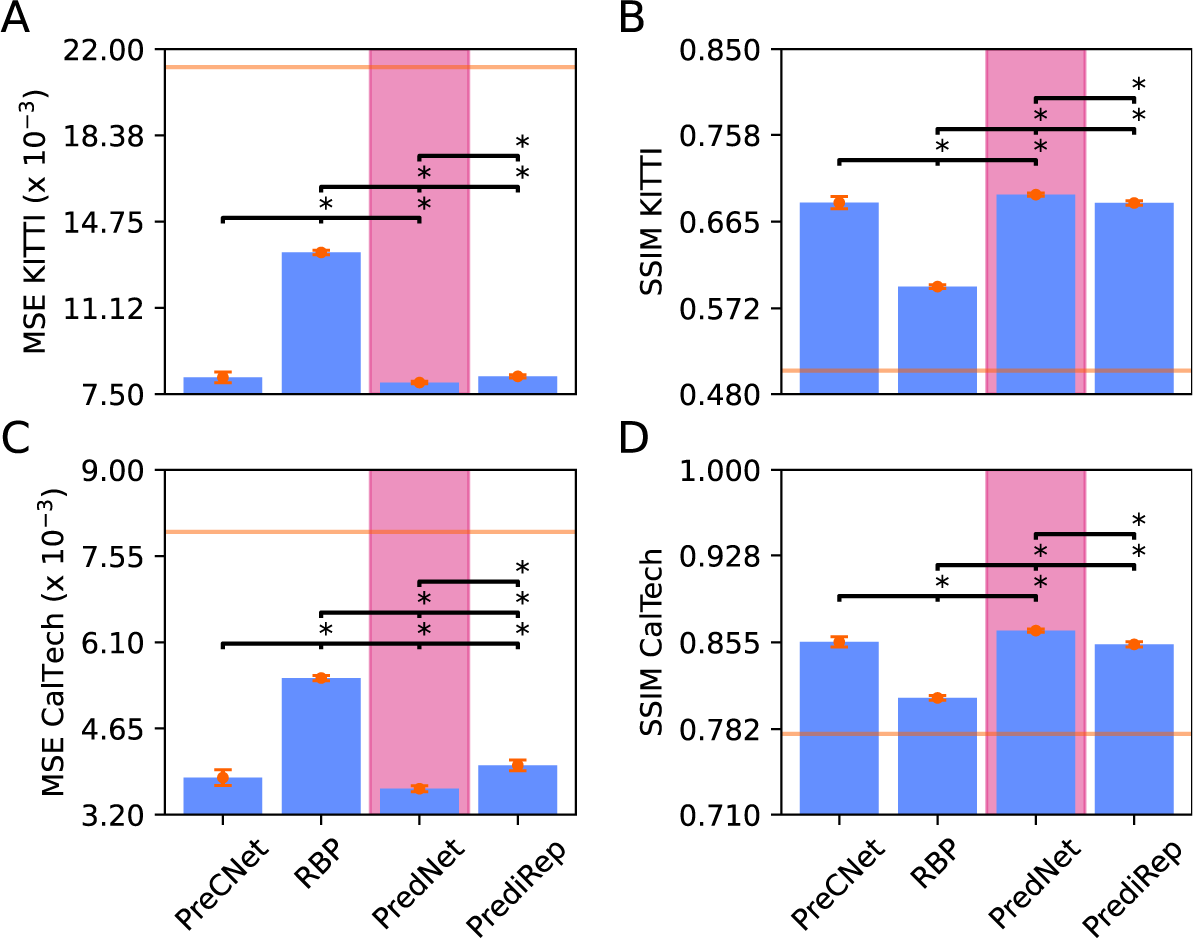
Performances of networks trained with Zero loss and a Tanh activation function except for PrediRep. This figure depicts the mean performances and distributions of PreCNet, RBP, PredNet and PrediRep for A) MSE on the KITTI dataset, B) SSIM for the KITTI dataset, C) MSE for the CalTech Pedestrian dataset and D) SSIM for the CalTech Pedestrian Dataset. All networks shown here used a Tanh as the activation function for their recurrent units except for PrediRep which used a ReLU. The pink color highlights the best performing model for each metric. The orange error bars depict the standard deviation of the performances of the networks. The horizontal solid orange lines indicate the performances achieved for each metric by simply copying the last frame. The asterisks above the black lines depict if the network on the left, from which the line originates, had a significantly different performance on the metric compared to the model above which the asterisk is placed. Statistics are based on Welch’s t-test with Bonferroni correction.

Figure 3 again presents the performances of the networks trained with Zero loss. However, in this case, all networks used ReLU as the activation function in their recurrent units compared to only PrediRep in Figure 2. PredNetZero emerged as the best performing network across multiple metrics, displaying superior results in all comparisons, apart from the MSE result for KITTI, where it was matched by PrediRepZero. PrediRepZero was the second-best performing network. It performed significantly better than PreCNetZero on the KITTI dataset while matching it on the CalTech Pedestrian dataset. It outperformed RBPZero across all metrics. In terms of the variability in the performances PrediRepZero, PredNetZero, and RBPZero exhibited consistent performances, with minimal variations. In contrast, PreCNetZero displayed a wide range of performances.

**Figure 3:**
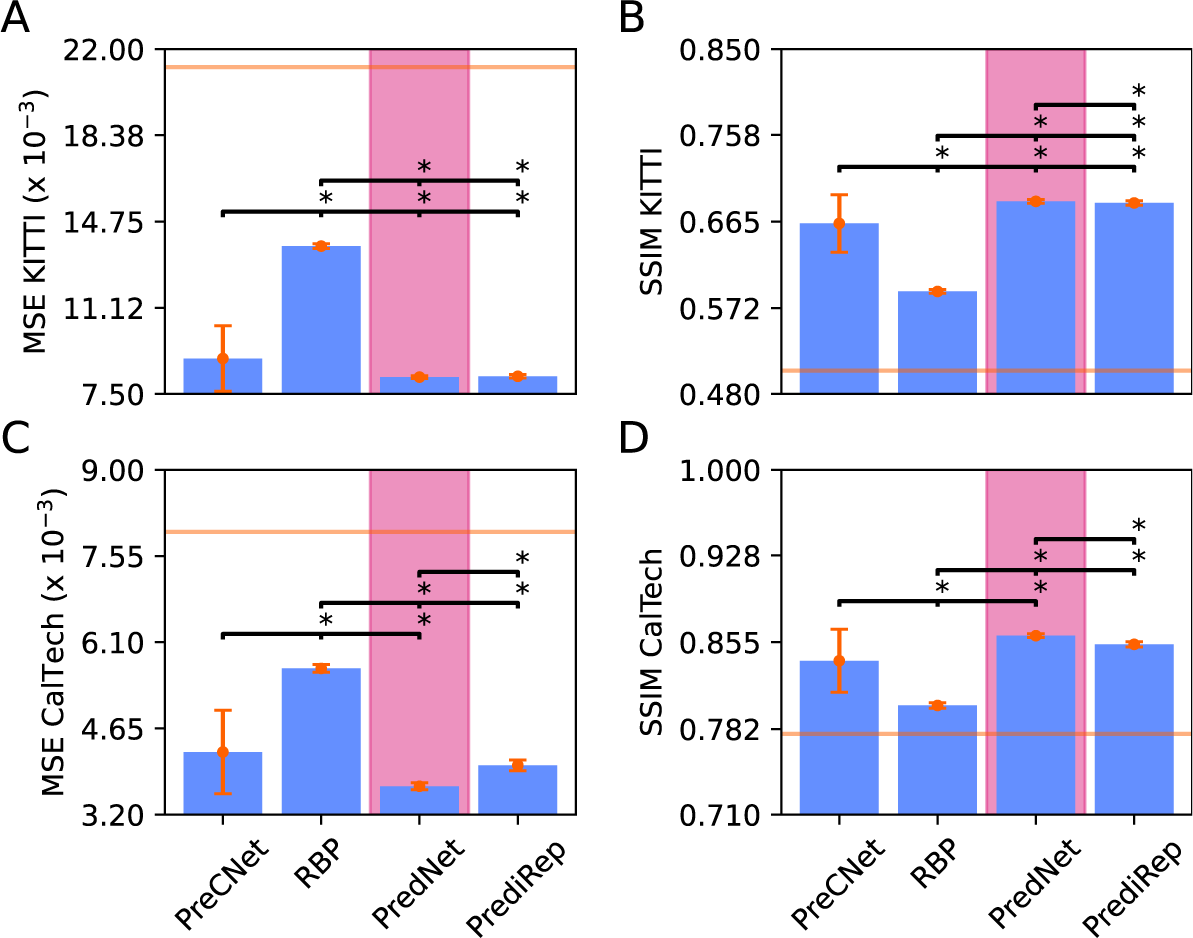
Performances of networks trained with Zero loss and a ReLU activation function. This figure depicts the mean performances and distributions of PreCNet, RBP, PredNet and PrediRep for A) MSE on the KITTI dataset, B) SSIM for the KITTI dataset, C) MSE for the CalTech Pedestrian dataset and D) SSIM for the CalTech Pedestrian Dataset. All networks here used a ReLU non-linearity as the activation function for their recurrent units. The pink color highlights the best performing model for each metric. The orange error bars depict the standard deviation of the performances of the networks. The horizontal solid orange lines indicate the performances achieved for each metric by simply copying the last frame. The asterisks above the black lines depict if the network on the left, from which the line originates, had a significantly different performance on the metric compared to the model above which the asterisk is placed. Statistics are based on Welch’s t-test with Bonferroni correction.

Figure 4 depicts the performances of the networks trained with All loss. In contrast to the networks trained with Zero loss, there was a substantial drop in performance, as indicated by the rise in the MSE and drop in the SSIM scores. This decrease in performance was also accompanied by an increase in performance variability. For the networks trained with All loss, PrediRepAll emerged as the best performing network, this time having significantly outperformed all networks on the KITTI dataset. Moreover, for both metrics on the CalTech Pedestrian dataset, PrediRepAll outperformed PreCNetAll and RBPAll by a significant margin. However, while performing better than PredNetAll in terms of SSIM performance on the CalTech Pedestrian dataset, it underperformed it for the MSE metric.

**Figure 4:**
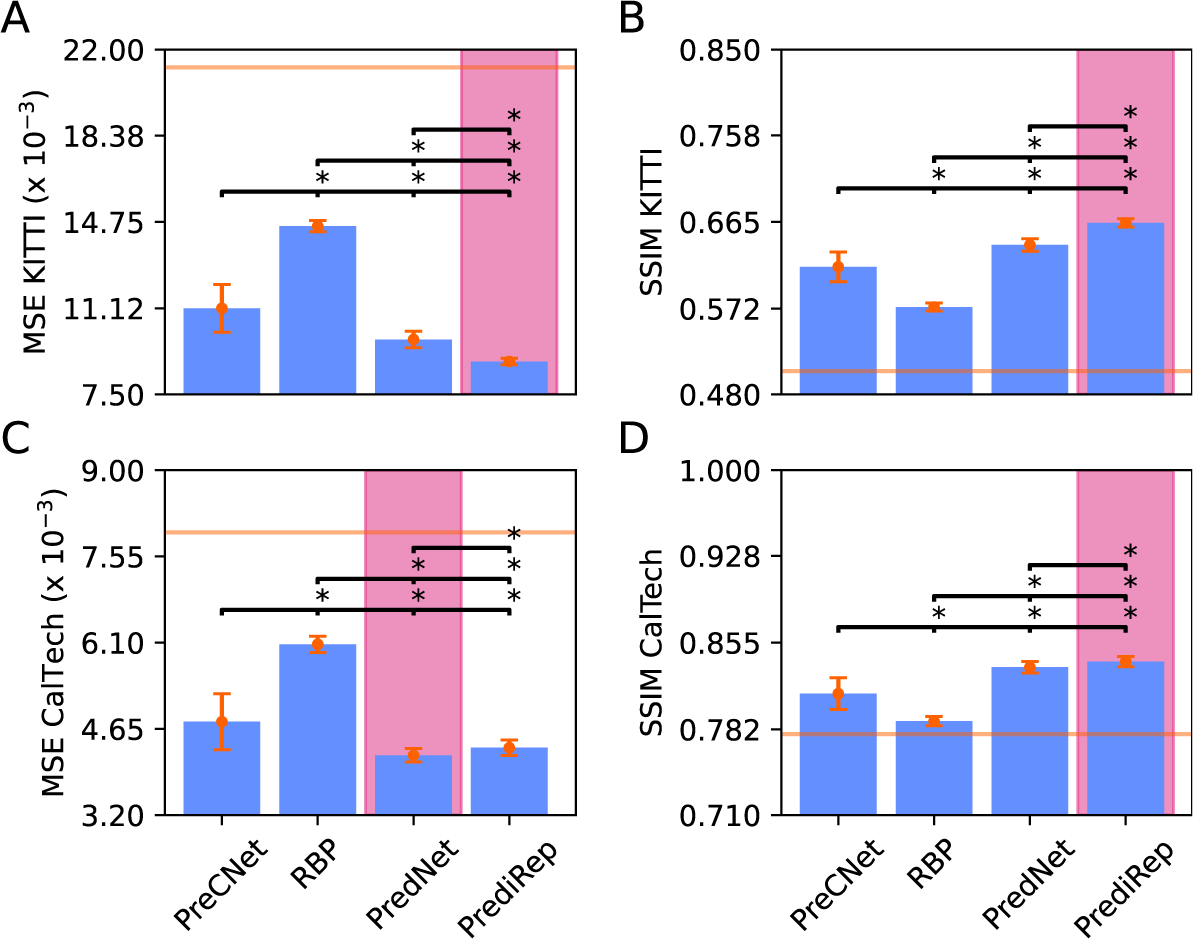
Performances of networks trained with All loss and a ReLU activation function. The description is the same as for Figure 3. This figure depicts the mean performances and distributions of PreCNet, RBP, PredNet and PrediRep for A) MSE on the KITTI dataset, B) SSIM for the KITTI dataset, C) MSE for the CalTech Pedestrian dataset and D) SSIM for the CalTech Pedestrian Dataset. All networks here used a ReLU non-linearity as the activation function for their recurrent units. The pink color highlights the best performing model for each metric. The orange error bars depict the standard deviation of the performances of the networks. The horizontal solid orange lines indicate the performances achieved for each metric by simply copying the last frame. The asterisks above the black lines depict if the network on the left, from which the line originates, had a significantly different performance on the metric compared to the model above which the asterisk is placed. Statistics are based on Welch’s t-test with Bonferroni correction.

Figure 5 presents the performances of the networks trained with Equal loss. Similarly to the transition from Zero to All loss, the switch from All to Equal loss resulted in performance declines and slight increases in performance variabilities among the networks. As for the networks trained with the All loss, PrediRepEqual showed, by a wide margin, the best performance on the KITTI datasets. In the case of the CalTech Pedestrian dataset, the performance of networks matched the level expected by simply copying the last frame for the SSIM metric. For the MSE metric, PrediRepEqual exhibited superior performance by outperforming both PredNetEqual and RBPEqual, while matching PreCNetEqual.

**Figure 5:**
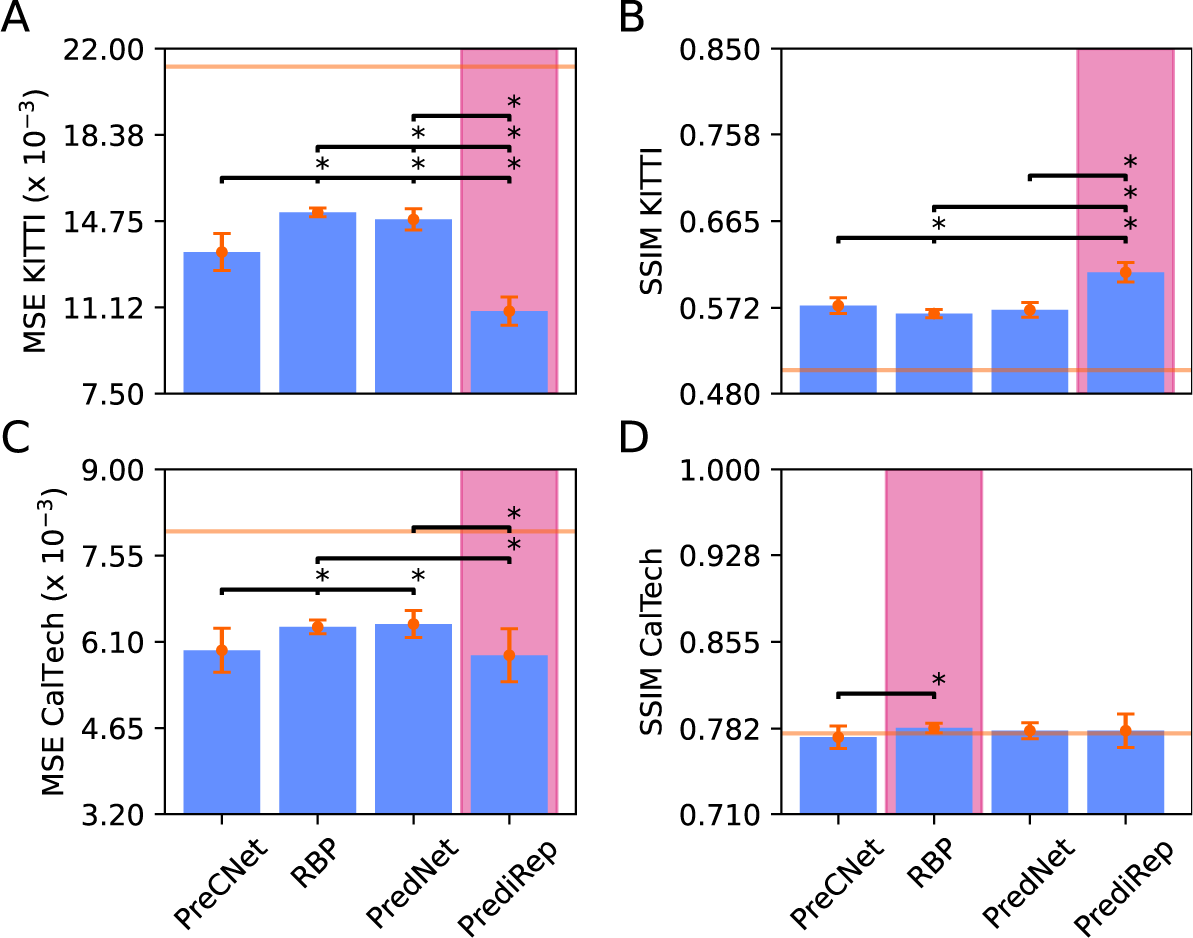
Performances of networks trained with Equal loss and a ReLU activation function. The description is the same as for Figure 3. This figure depicts the mean performances and distributions of PreCNet, RBP, PredNet and PrediRep for A) MSE on the KITTI dataset, B) SSIM for the KITTI dataset, C) MSE for the CalTech Pedestrian dataset and D) SSIM for the CalTech Pedestrian Dataset. All networks here used a ReLU non-linearity as the activation function for their recurrent units. The pink color highlights the best performing model for each metric. The orange error bars depict the standard deviation of the performances of the networks. The horizontal solid orange lines indicate the performances achieved for each metric by simply copying the last frame. The asterisks above the black lines depict if the network on the left, from which the line originates, had a significantly different performance on the metric compared to the model above which the asterisk is placed. Statistics are based on Welch’s t-test with Bonferroni correction.

### 3.2. Representation units should make accurate predictions of the targets

According to the principles of hPC, R neurons aim to make precise predictions of lower-level activity. In the context of the networks evaluated in this study, the accuracy of the predictions generated by the R units was assessed by calculating the correlations between the predictions and the targets for each level. This analysis was conducted for the final frame of all sequences within the KITTI test set. Figures 6A-C provide a visualization of these results. In all instances of the Zero networks, as seen in panel 6A, the R units did not exhibit high accuracy in predicting targets beyond level 0. PrediRepZero was, from all networks trained with Zero loss, the network with the most robust correlations at higher levels. PredNetAll, displayed in panel 6B, did not show an increase in correlation for higher levels when compared to PredNetZero. Furthermore, both PredNetEqual and RBPEqual did not have correlation values for higher levels as either the targets or predictions for those levels were empty and are therefore not depicted in panel 6C. This also applied to RBPAll for levels 2 and 3. In contrast, PrediRep and PreCNet networks trained with All and Equal loss consistently demonstrated accurate predictions of the targets across all levels. PrediRep exhibited slightly superior performance for both loss types.

**Figure 6:**
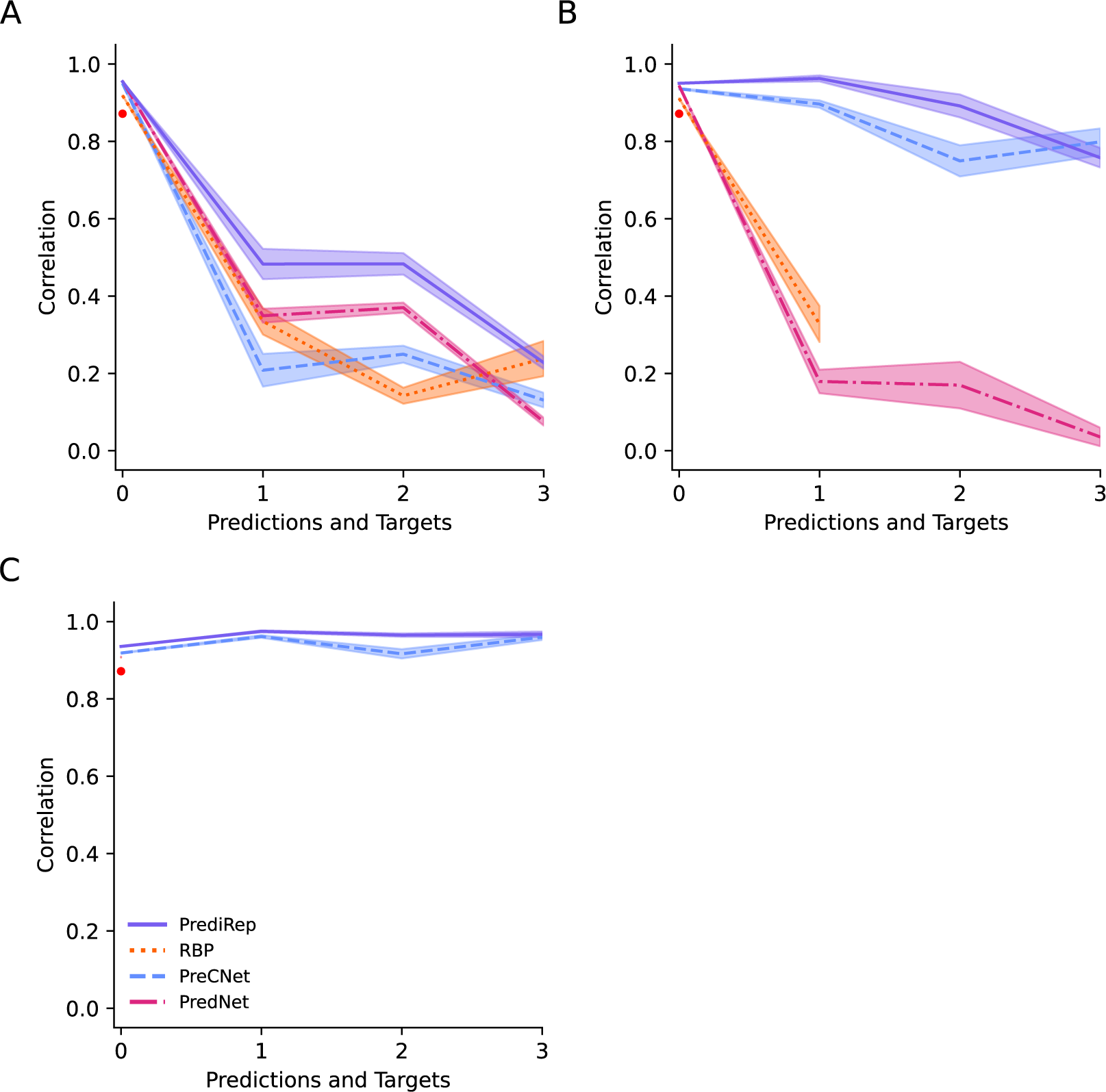
Correlations between predictions and targets. This figure shows the mean correlations between the predictions generated by the representation units and their targets for networks trained with A) Zero loss, B) All loss, and C) Equal loss. The correlations are averaged across all instantiations of the different combinations of network and loss types. Correlations are given for each of the four levels of the networks with predictions. Some combinations lacked correlation values for certain levels as the predictions or the targets were empty for all instances of that combination. The small red dots depict the results of a baseline model which simply correlates the last frame with the previous one. The distributions around the means are based on the standard error of the means.

### 3.3. Representation units and their predictions should not fall silent

Figures 6A-C revealed that specific combinations of network and loss type result in empty predictions or targets across all instances of that combination. However, adherence to the principles of hPC necessitates that, in the presence of a stimulus, the R neurons and their predictions must remain active. To assess the compliance of these instantiations with this expectation, an examination of the R units and their predictions for the final frame was conducted across all unique sequences in the KITTI test set. These findings are presented in Table 3. In the case of networks trained with Zero loss, almost all instances, with a few exceptions of RBPZero, consistently produced non-zero representations and predictions. Conversely, for runs trained with All and Equal losses the results present a striking contrast. None of the instances of the networks of RBPAll, RBPEqual and PredNetEqual displayed the expected behavior. PreCNetEqual, PreCNetAll and PredNetAll exhibited only a limited number of instances that met this expectation. On the other hand, PrediRepAll and PrediRepEqual consistently generated non-zero targets and predictions for the networks.

**Table 3:** Number of Instantiations of a Network and Loss Size Combination Where Representation Units and Their Predictions Were Active.

| Network | Zero Loss | All Loss | Equal Loss |
| --- | --- | --- | --- |
| PreCNet | 30 | 11 | 4 |
| RBP | 27 | 0 | 0 |
| PredNet | 30 | 4 | 0 |
| PrediRep | 30 | 29 | 25 |

### 3.4. Activities in representation units and their predictions should correlate with sensory input

In hierarchical generative models, a central argument posits that distinct levels of the cortex process the same underlying information but at varying levels of abstraction. Cortical areas at the lower levels of the hierarchy primarily predict low-level features. Concurrently, cortical areas at higher levels become increasingly involved in the processing of more abstract and complex aspects of the same information and therefore predict more high-level features (Mumford, 1992; Friston, 2005; Clark, 2013). This concept suggests that the closer a level is to the sensory input, the more correlated it should be with it. Furthermore, this correlation should exhibit a monotonically decreasing trend when moving up through the hierarchical levels. In the context of hPC this implies that R neurons should display correlation with the sensory input, and this correlation should diminish across levels. Moreover, since the predictions emanating from the R units should reflect the information contained in their counterparts of the level below, it implies that the predictions should also display high correlations with the sensory input.

To investigate whether this expected behavior occurred in the network instantiations, the correlations between the activities of the R units of each level and their predictions with the frames were measured. In the case of the RBP network, the frames selected for the correlation depended on the specific level being analyzed due to the inherent lag effect between levels of the network. For the other networks, the final frame of a sequence was consistently used. Depending on the dimensions of the R or Ậ unit activity, the frames were downsampled using stacked (2 x 2) max pooling operations. This correlation analysis encompassed all sequences within the KITTI test set.

Figures 7A-C shows that in most network and loss type combinations, the R units showed the expected monotonic decrease in correlation with the frames when ascending through the hierarchy. One combination that deviated from this pattern was PreCNetEqual which demonstrated a significant increase in correlation from level 1 to level 2 as can be seen in Figure 7C. PrediRep stood out clearly from the other networks regardless of the loss type as it consistently exhibited notably higher correlations between the R units and the frames across levels. Moreover, there was a substantial increase in the correlation from PrediRepZero to PrediRepAll, followed by a smaller increase to PrediRepEqual.

**Figure 7:**
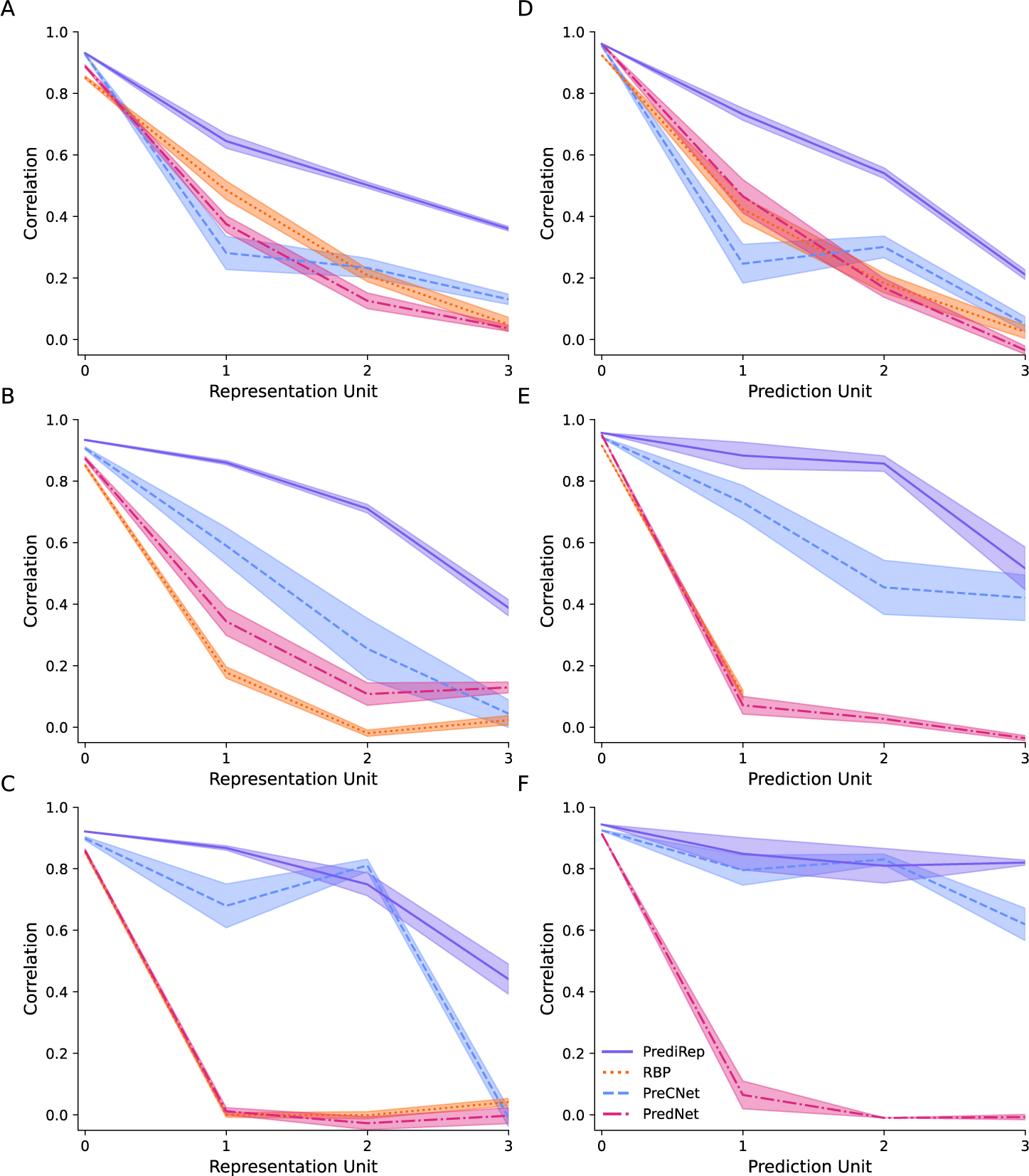
Correlations between representation units and frames and their predictions and frames. This figure represents the mean correlations between the representation units and frames for networks trained with A) Zero loss, B) All loss, and C) Equal loss and the predictions of representation units and frames for networks trained with D) Zero loss, E) All loss, and F) Equal loss. Correlations are shown for each representation unit and their prediction. The distributions around the means are based on the standard error of the means.

The results depicted in Figures 7D-F were similar to those found in Figures 7A-C. Once again, most network and loss type combinations showed a monotonic decrease in correlation across levels. PreCNetEqual again stood out, in Figure 7F, with an increase in correlation from level 1 to 2. Furthermore, the correlations of PrediRep consistently surpassed those of all other networks for every loss type. It is however important to note that PrediRepEqual did not show the expected monotonic decrease with levels. There was a slight increase in the correlation of the prediction in level 2 compared to that in level 1. Nonetheless, PrediRepAll maintained the anticipated monotonic decrease in correlation.

## 4. Discussion

The results of our study can be interpreted with a dual focus, either emphasizing the functional alignment with hPC or emphasizing computational performance. Viewed from the hPC perspective, our findings underscore the crucial role of the loss type employed when training the networks. It is evident that the networks under consideration cannot be adequately evaluated in isolation from the chosen loss type. Importantly, the results decidedly point to PrediRepAll as the most promising model of hPC. Several key observations and analyses support this conclusion. Firstly, Figures 6A-C revealed that certain network and loss type combinations, particularly those trained with Zero loss, do not serve as suitable models of hPC. This is attributed to the fact that the instantiations of these models form R units that do not generate accurate predictions of the targets at higher levels. Adding to this issue, some combinations do not develop R units that exhibit any activity whatsoever. Specifically, not a single instance of RBPAll, RBPEqual, or PredNetEqual showed R units that made active predictions, as evidenced in Table 3. In contrast, the majority of PrediRep instances trained with either All or Equal loss exhibited R units that were both active and made accurate predictions of their targets. Networks that did produce some instances with precise higher-level predictions, such as PreCNetEqual and PreCNetAll, did so to a much lesser extent compared to their respective PrediRep counterparts. Additionally, Figures 7A-F highlighted that the R and Ậ units of the PreCNet networks exhibited a lower correlation with the frames compared to those of the PrediRep networks. Between PrediRepAll and PrediRepEqual the former emerges as more favorable due to the balanced results for various functional aspects of hPC. Therefore, the work presented here reveals PrediRepAll as a robust candidate for modeling hPC functionality.

From a computational performance standpoint, the results of our investigations align with the findings of Lotter et al. (2016), demonstrating that networks trained with Zero loss consistently yield superior results compared to other loss types. Moreover, using a Tanh non-linearity as the activation function for the ConvLSTM units also leads to a small boost in performance as found in Lotter et al. (2020). When evaluating computational performance across all network types, PredNetZeroTanh emerged as the top-performing model. Intriguingly, the compact PrediRepZero, with only half the number of the trainable parameters compared to the other networks and without using ConvLSTM units, consistently achieved competitive performance in comparison to its larger counterparts. Previously, Straka et al. (2023) demonstrated that PreCNet outperformed PredNet when the two networks shared similar parameter numbers. However, it is important to note that their PreCNet architecture featured a substantially higher number of filters in the lowest R unit compared to their PredNet implementation. In contrast, our current findings indicate that, when PredNet and PreCNet possess an identical number of filters, PredNet outperforms PreCNet. This suggests that augmenting the number of filters in the R units, especially the lowest one, enhances a network’s ability to predict the next frame. The unique advantage of PrediRep lies in its lightweight design, enabling the incorporation of more filters in the R units while maintaining fewer overall trainable parameters compared to the other networks.

Table 3 highlights that, in the case of All and Equal losses, both PredNet and RBP instantiations exhibit inconsistent formation of active representations and predictions. In contrast, PrediRep and PreCNet consistently demonstrate the desired behavior. The delineating factor between these network types, presumably contributing to the differences in consistency, lies in the algorithms of PredNet and the RBP. Regardless of the network type, the target entering the error units of the network at the lowest level is always the unmodified sensory input. However, in higher-levels for PredNet and RBP, the algorithms incorporate mechanisms by which the information in the representation units (RBP) or error units (PredNet) are modified via learnable weights before becoming the targets for the next-level. This mechanism of altering information entering higher-level error units coupled with the objective of minimizing the error unit activity for all levels under the All and Equal loss configurations, may induce PredNet and RBP to learn to nullify the activity entering these units. This phenomenon is absent in PrediRep and PreCNet as they do not contain a mechanism by which the information entering the error units can be modified by learnable weights.

PrediRepAll demonstrates a robust functional congruence with hPC while maintaining a neutral stance on the precise implementation of hPC in the cortex, specifically on the role of feedback connections. Previous research indicates that inter-cortical feedback connections originate from deep layers and target both the superficial and deep layers of the receiving area (Felleman and Van Essen, 1991). Newer investigations further propose the possible existence of a second feedback stream originating in superficial layers (Markov et al., 2014; Vezoli et al., 2021). Given these various possibilities of feedback connections, models devised to explain the implementation of hPC vary in their assertions about the connections responsible for specific functional aspects (Bastos et al., 2012; Shipp, 2016), such as the identity of the feedback connection conveying predictions. Consequently, PrediRepAll is presented as an abstract model of hPC, encompassing necessary units and connections posited to exist in hPC without making explicit claims about the specific cortical connectivity.

Recent advancements in the pursuit of enhancing the biological plausibility of predictive coding have led to the emergence of dendritic predictive coding (dPC; Mikulasch et al., 2023; Tang et al., 2023; Millidge et al., 2023). Within the framework of dPC, a shift is made by relocating error computations from explicit error neurons to the dendritic compartments of pyramidal neurons. In this context, prediction errors are postulated to be reflected in the difference in membrane potential between the soma and the apical dendrites of these neurons. PrediRep, as stated before, is an abstract model of hPC and does not impose the specific requirements that the units in the R and E units must represent separate neurons. Therefore, it readily accommodates alignment with dPC by a straightforward redefinition of these separate units, combining them into a single unit where the representation component corresponds to the soma, and the error component to the apical dendrite.

While our study has successfully explored the functional alignment of PrediRepAll with hPC, it is important to acknowledge the limited scope of our investigation. As a neuroscientific theory, hPC encompasses a multitude of cortical phenomena which have not been considered here. For instance, hPC has been utilized to explain extra-classical receptive field effects such as end-stopping (Hubel and Wiesel, 1968; Rao and Ballard, 1999). Furthermore, hPC has been implicated as a potential mechanism behind the role of predictive feedback involved in when viewing occluded images (Muckli et al., 2015; Morgan et al., 2019) and illusions (Kok et al., 2016). Moreover, considering that hPC posits distinct laminar locations for R and E neurons, it has been applied in attempts to explain cortical layer data derived from laminar fMRI studies (Muckli et al., 2015; Kok et al., 2016; Aitken et al., 2020; Haarsma et al., 2022). Therefore, there remains a wide spectrum of opportunities for further scrutinizing the alignment of PrediRepAll with hPC.

Our findings clearly show that minor deviations from the principles of a neuroscientific theory may lead to major deviations from the functional implications of that theory. We introduce PrediRep, a novel recurrent convolutional neural network that reproduces functional features of cortical processing as proposed by predictive coding while maintaining competitive performance on machine vision tasks. This renders PrediRep well suited for investigating cortical phenomena within the context of hPC.

## 5. Acknowledgments

This study has received funding from the European Union’s Horizon 2020 Framework Programme for Research and Innovation under the Specific Grant Agreement 945539 (Human Brain Project SGA3). We would like to thank Prof. dr. Federico De Martino for the helpful discussions on predictive coding.

## 6. Declaration of Generative AI and AI-assisted technologies in the writing process

During the preparation of this work the author(s) used ChatGPT3.5 in order to improve readability and language. After using this tool/service, the author(s) reviewed and edited the content as needed and take(s) full responsibility for the content of the publication.

## Footnotes

1 https://github.com/ccnmaastricht/predirep

## References

Aitken, F., Menelaou, G., Warrington, O., Koolschijn, R. S., Corbin, N., Callaghan, M. F., and Kok, P. (2020). Prior expectations evoke stimulus-specific activity in the deep layers of the primary visual cortex. PLOS Biology, 18(12):e3001023. Number: 12.

Bastos, A., Usrey, W., Adams, R., Mangun, G., Fries, P., and Friston, K. (2012). Canonical Microcircuits for Predictive Coding. Neuron, 76(4):695– 711. Number: 4.

Clark, A. (2013). Whatever next? Predictive brains, situated agents, and the future of cognitive science. Behavioral and Brain Sciences, 36(3):181–204. Number: 3.

Dollar, P., Wojek, C., Schiele, B., and Perona, P. (2009). Pedestrian detection: A benchmark. In 2009 IEEE Conference on Computer Vision and Pattern Recognition, pages 304–311, Miami, FL. IEEE.

Felleman, D. J. and Van Essen, D. C. (1991). Distributed Hierarchical Processing in the Primate Cerebral Cortex. Cerebral Cortex, 1(1):1–47. Number: 1.

Finn, C., Goodfellow, I., and Levine, S. (2016). Unsupervised Learning for Physical Interaction through Video Prediction. In Proceedings of the 30th International Conference on Neural Information Processing Systems, NIPS’16, pages 64–72, Red Hook, NY, USA. Curran Associates Inc. event-place: Barcelona, Spain.

Friston, K. (2003). Learning and inference in the brain. Neural Networks, 16(9):1325–1352. Number: 9.

Friston, K. (2005). A theory of cortical responses. Philosophical Transactions of the Royal Society B: Biological Sciences, 360(1456):815–836. Number: 1456.

Geiger, A., Lenz, P., Stiller, C., and Urtasun, R. (2013). Vision meets robotics: The KITTI dataset. The International Journal of Robotics Research, 32(11):1231–1237. Number: 11.

Goroshin, R., Mathieu, M. F., and LeCun, Y. (2015). Learning to Linearize Under Uncertainty. In Cortes, C., Lawrence, N., Lee, D., Sugiyama, M., and Garnett, R., editors, Advances in Neural Information Processing Systems, volume 28. Curran Associates, Inc.

Haarsma, J., Kok, P., and Browning, M. (2022). The promise of layer-specific neuroimaging for testing predictive coding theories of psychosis. Schizophrenia Research, 245:68–76.

Hosseini, M. and Maida, A. (2020). Hierarchical Predictive Coding Models in a Deep-Learning Framework. Issue: arXiv:2005.03230 arXiv:2005.03230 [cs].

Huang, Y. and Rao, R. P. N. (2011). Predictive coding. WIREs Cognitive Science, 2(5):580–593. Number: 5.

Hubel, D. H. and Wiesel, T. N. (1968). Receptive fields and functional architecture of monkey striate cortex. The Journal of Physiology, 195(1):215– 243.

Jiang, L. P. and Rao, R. P. (2022). Predictive Coding Theories of Cortical Function. In Oxford Research Encyclopedia of Neuroscience. Oxford University Press. Book Authors: n31.

Kingma, D. P. and Ba, J. (2014). Adam: A Method for Stochastic Optimization. Publisher: arXiv Version Number: 9.

Kirubeswaran, O. and Storrs, K. R. (2023). Inconsistent illusory motion in predictive coding deep neural networks. Vision Research, 206:108195.

Kok, P., Bains, L., van Mourik, T., Norris, D., and de Lange, F. (2016). Selective Activation of the Deep Layers of the Human Primary Visual Cortex by Top-Down Feedback. Current Biology, 26(3):371–376. Number: 3.

Lee, T. S. and Mumford, D. (2003). Hierarchical Bayesian inference in the visual cortex. Journal of the Optical Society of America A, 20(7):1434. Number: 7.

Lotter, W., Kreiman, G., and Cox, D. (2016). Deep Predictive Coding Networks for Video Prediction and Unsupervised Learning. Publisher: arXiv Version Number: 5.

Lotter, W., Kreiman, G., and Cox, D. (2020). A neural network trained for prediction mimics diverse features of biological neurons and perception. Nature Machine Intelligence, 2(4):210–219. Number: 4.

Markov, N. T., Vezoli, J., Chameau, P., Falchier, A., Quilodran, R., Huissoud, C., Lamy, C., Misery, P., Giroud, P., Ullman, S., Barone, P., Dehay, C., Knoblauch, K., and Kennedy, H. (2014). Anatomy of hierarchy: Feedforward and feedback pathways in macaque visual cortex. Journal of Comparative Neurology, 522(1):225–259.

Mathieu, M., Couprie, C., and LeCun, Y. (2015). Deep multi-scale video prediction beyond mean square error. Publisher: arXiv Version Number: 6.

Mikulasch, F. A., Rudelt, L., Wibral, M., and Priesemann, V. (2023). Where is the error? Hierarchical predictive coding through dendritic error computation. Trends in Neurosciences, 46(1):45–59. Number: 1.

Millidge, B., Tang, M., Osanlouy, M., and Bogacz, R. (2023). Predictive Coding Networks for Temporal Prediction. preprint, Neuroscience.

Morgan, A. T., Petro, L. S., and Muckli, L. (2019). Scene Representations Conveyed by Cortical Feedback to Early Visual Cortex Can Be Described by Line Drawings. The Journal of Neuroscience, 39(47):9410–9423. Number: 47.

Muckli, L., De Martino, F., Vizioli, L., Petro, L., Smith, F., Ugurbil, K., Goebel, R., and Yacoub, E. (2015). Contextual Feedback to Superficial Layers of V1. Current Biology, 25(20):2690–2695. Number: 20.

Mumford, D. (1992). On the computational architecture of the neocortex: II The role of cortico-cortical loops. Biological Cybernetics, 66(3):241–251.

O’Reilly, R. C., Wyatte, D., and Rohrlich, J. (2014). Learning Through Time in the Thalamocortical Loops. Publisher: arXiv Version Number: 1.

Palm, R. B. (2012). Prediction as a candidate for learning deep hierarchical models of data.

Patraucean, V., Handa, A., and Cipolla, R. (2015). Spatio-temporal video autoencoder with differentiable memory. Publisher: arXiv Version Number: 5.

Rao, R. P. N. and Ballard, D. H. (1998). Development of localized oriented receptive fields by learning a translation-invariant code for natural images. Network: Computation in Neural Systems, 9(2):219–234. Number: 2.

Rao, R. P. N. and Ballard, D. H. (1999). Predictive coding in the visual cortex: a functional interpretation of some extra-classical receptive-field effects. Nature Neuroscience, 2(1):79–87.

Shi, X., Chen, Z., Wang, H., Yeung, D.-Y., Wong, W.-k., and Woo, W.-c. (2015). Convolutional LSTM Network: A Machine Learning Approach for Precipitation Nowcasting. Publisher: arXiv Version Number: 2.

Shipp, S. (2016). Neural Elements for Predictive Coding. Frontiers in Psychology, 7.

Softky, W. (1995). Unsupervised Pixel-prediction. In Touretzky, D., Mozer, M. C., and Hasselmo, M., editors, Advances in Neural Information Processing Systems, volume 8. MIT Press.

Spratling, M. (2017). A review of predictive coding algorithms. Brain and Cognition, 112:92–97.

Srivastava, N., Mansimov, E., and Salakhutdinov, R. (2015). Unsupervised Learning of Video Representations using LSTMs. Publisher: arXiv Version Number: 3.

Straka, Z., Svoboda, T., and Hoffmann, M. (2023). PreCNet: Next-Frame Video Prediction Based on Predictive Coding. IEEE Transactions on Neural Networks and Learning Systems, pages 1–15.

Tang, M., Salvatori, T., Millidge, B., Song, Y., Lukasiewicz, T., and Bogacz, R. (2023). Recurrent predictive coding models for associative memory employing covariance learning. PLOS Computational Biology, 19(4):e1010719. Number: 4.

Vezoli, J., Magrou, L., Goebel, R., Wang, X.-J., Knoblauch, K., Vinck, M., and Kennedy, H. (2021). Cortical hierarchy, dual counterstream architecture and the importance of top-down generative networks. NeuroImage, 225:117479.

Vondrick, C., Pirsiavash, H., and Torralba, A. (2016). Generating Videos with Scene Dynamics. In Lee, D., Sugiyama, M., Luxburg, U., Guyon, I., and Garnett, R., editors, Advances in Neural Information Processing Systems, volume 29. Curran Associates, Inc.

Wang, Z., Bovik, A., Sheikh, H., and Simoncelli, E. (2004). Image Quality Assessment: From Error Visibility to Structural Similarity. IEEE Transactions on Image Processing, 13(4):600–612. Number: 4.

Watanabe, E., Kitaoka, A., Sakamoto, K., Yasugi, M., and Tanaka, K. (2018). Illusory Motion Reproduced by Deep Neural Networks Trained for Prediction. Frontiers in Psychology, 9:345.

Werbos, P. (1990). Backpropagation through time: what it does and how to do it. Proceedings of the IEEE, 78(10):1550–1560.

